# Auditory-Visual Speech Behaviors are Resilient to Left pSTS Damage

**DOI:** 10.1101/2020.09.26.314799

**Authors:** David Brang, John Plass, Sofia Kakaizada, Shawn L. Hervey-Jumper

**Affiliations:** Department of Psychology, University of Michigan, Ann Arbor, MI 48109; Department of Neurological Surgery, University of California, San Francisco, San Francisco, CA, 94115

**Keywords:** Multisensory, Audiovisual, Lesion, Language, Tumor, Recovery

## Abstract

The ability to understand spoken language is essential for social, vocational, and emotional health, but can be disrupted by environmental noise, injury, or hearing loss. These auditory deficits can be ameliorated by visual speech signals that convey redundant or supplemental speech information, but the brain regions critically responsible for these audiovisual (AV) interactions remain poorly understood. Previous TMS and lesion-mapping studies suggest that the left posterior superior temporal sulcus (pSTS) is causally implicated in producing the McGurk effect, an AV illusion in which auditory and visual speech are perceptually “fused.” However, previous research suggests that the McGurk effect is neurally and behaviorally dissociable from other visual effects on speech perception and, therefore, may not provide a generalizable index of AV interactions in speech perception more broadly. To examine whether the left pSTS is critically responsible for AV speech integration more broadly, we measured the strength of the McGurk effect, AV facilitation effects, and AV conflict effects longitudinally over 2 years in patients undergoing surgery for intrinsic tumors in the left pSTS (*n* = 2) or frontal lobes (control; *n* = 14). Results demonstrated that left pSTS lesions impaired experience of the McGurk effect, but did not uniformly reduce visual influences on speech perception. Additionally, when multisensory behaviors were affected by a lesion, AV speech perception abilities could recover over time. Our results suggest a causal dissociation between perceptual benefits produced by congruent AV speech and perceptual modulations produced by incongruent AV speech (the McGurk effect).These data are consistent with models proposing that that the pSTS is only one of multiple critical areas necessary for AV speech interactions.

## INTRODUCTION

Speech perception is essential for social, vocational, and emotional health in hearing individuals (Blumstein, 1994; Ziegler, Pech-Georgel, George, Alario, & Lorenzi, 2005). However, auditory speech signals can be degraded by environmental noise (e.g., a crowded party), sensory/neural damage, or age-related hearing loss, necessitating supplemental processes to enable fast and accurate recovery of speech information (Irwin & DiBlasi, 2017; Puschmann et al., 2019). The principle way this occurs in healthy individuals is through the use of visual speech cues (Grant & Seitz, 2000; Sumby & Pollack, 1954). However, the neural mechanisms that are critically required for visual facilitation of speech perception remain poorly understood.

Studies of the McGurk Effect -- in which an auditory phoneme such as /ba/ paired with the image of a speaker saying /ga/ results in the fused percept of /da/ (McGurk & MacDonald, 1976) -- have led to a focus on the left posterior superior temporal sulcus (pSTS) as the putative region responsible for enabling multisensory speech perception (Beauchamp, Nath, & Pasalar, 2010). For example, individual differences in the strength of the McGurk effect are correlated with fMRI activity in the pSTS during the McGurk effect (Nath & Beauchamp, 2012) and either inhibitory transcranial magnetic stimulation (TMS) or damage following a stroke in this region reduces the strength of the McGurk effect (Beauchamp et al., 2010; Hickok et al., 2018).

However, several studies have recently questioned the generalizability of models based on the McGurk effect in explaining visual effects on speech perception more broadly (Alsius, Paré, & Munhall, 2018; Ganesan et al., 2020). Evidence from behavioral, electrophysiological, and neuroimaging studies suggests that visual facilitation of speech perception (e.g., improved detection in noise) likely relies on different neural mechanisms than visual modulations of speech perception (e.g., the McGurk effect). For example, while the McGurk effect requires explicit recognition of audiovisual (AV) speech signals as speech, audiovisual facilitations of speech perception do not (K. Eskelund, J. Tuomainen, & T. Andersen, 2011; Munhall, MacDonald, Byrne, & Johnsrude, 2009; Palmer & Ramsey, 2012; J Plass, Guzman-Martinez, Ortega, Grabowecky, & Suzuki, 2014), suggesting that the McGurk effect may rely on a speech-specific mechanism that is distinct from other more general mechanisms for crossmodal facilitation.

Consistent with this interpretation, neural responses associated with visual facilitation versus modulation of speech perception have been found to differ in their timing, topography, and spectral profiles. While visual facilitations of speech perception likely rely on early feedforward signals (delta-theta band, 2-7 Hz) that convey visually-derived temporal or acoustic information to auditory cortex, visual modulations of speech perception likely rely on later feedback (low beta band, 14-15 Hz) from higher-order pSTS mechanisms that integrate discrepant auditory and visual speech signals (Arnal, Morillon, Kell, & Giraud, 2009; Arnal, Wyart, & Giraud, 2011). Consistent with behavioral observations, early (~100 ms) effects associated with AV facilitation occur regardless of whether ambiguous stimuli are perceived as speech or non-speech, while later effects associated with AV incongruity (~200 ms) are only observed when stimuli are explicitly perceived as speech (Baart, Stekelenburg, & Vroomen, 2014; Stekelenburg & Vroomen, 2007). Taken together, these results suggest that AV facilitations of speech perception may rely on a different mechanism than the pSTS feedback mechanism associated with the McGurk effect. Because audio-visually congruent stimuli that produce perceptual facilitations are more ecologically valid than incongruent stimuli that produce the McGurk effect, examining the mechanisms subserving AV facilitations may be particularly important for understanding multisensory interactions in everyday speech perception.

Therefore, to understand whether the left pSTS is critically responsible for AV speech integration more broadly, it is necessary to examine its causal role in the integration of naturally congruent AV speech in addition to illusions such as the McGurk effect. Furthermore, to identify the functional and clinical implications of damage to this region, particularly as it compensates for receptive speech processes, it is necessary to examine how left pSTS damage effects AV speech perception over the time-course of treatment and subsequent recovery.

To address these issues, we examined AV speech perception longitudinally (~2 years) in 2 patients with intrinsic brain tumors affecting the left pSTS and compared their performance to 14 patients with comparable frontal lobe tumors. Results demonstrated that a lesion in the left pSTS did not always reduce visual influences on speech perception, but when multisensory behaviors were affected by a lesion, AV speech perception abilities could recover over time (presumably through compensatory neural plasticity). Furthermore, our results suggest a causal dissociation between perceptual benefits produced by congruent AV speech and perceptual modulations produced by incongruent AV speech (the McGurk effect).

## MATERIALS AND METHODS

### Participants

16 right-handed patients with an intrinsic brain tumor within either the left pSTS (*n* = 2) or the frontal lobe (*n* = 5 left frontal, *n* = 9 right frontal) were recruited from the University of California, San Francisco Medical Center. Patients ranged in age from 25 - 72 (mean = 46.1, SD = 15.3), including 7 female and 9 male individuals. These patients were selected from a larger ongoing study based the location of their tumor and as they had completed both a pre-operative testing session and at least one additional session at a minimum of 90 days following surgical resection of their tumor. Written consent was obtained from each participant according to the direction of the institutional review board at the University of California San Francisco. Patients participated as volunteers during the administration of other clinical procedures and received no financial compensation.

50 healthy undergraduate students (*n* = 44 right-handed) were recruited from the University of Michigan to provide normative data and operate as a non-clinical control group. Participants ranged in age from 17 - 22 (mean = 18.9, SD = 1.1), including 36 female and 14 male individuals. All participants gave informed consent prior to the experiment and were given course credit for their participation. This study was approved by the institutional review board at the University of Michigan.

### Method Details

#### Auditory-Visual Speech Paradigm

Patients were seated in a clinical testing room at the University of California San Francisco hospital. Auditory-visual stimuli were delivered via a laptop using PsychToolbox (Brainard, 1997; Pelli, 1997). Non-clinical control participants were seated in an experimental testing room and completed a matched paradigm on a desktop computer using free-field speakers.

The AV paradigm was designed to capture multiple elements of AV speech integration. Specifically, it involved AV movies from a prior study (Ross et al., 2007) in which a female speaker produces common monosyllabic words. From this stimulus set we selected 40 one-syllable words that each had one of four initial consonants: ‘b’, ‘f’, ‘g’, and ‘d’ (10 of each); phonemes in the second position of these words were additionally balanced across the four groups. Stimuli were recorded at 29.97 frames per second, trimmed to 1100 ms in length, and adjusted so the first consonantal burst of sound occurred at 500 ms during each video.

We designed the task conditions to orthogonally manipulate (1) the visual context present on a trial (no visual [auditory-alone], congruent AV, incongruent AV), and (2) the level of background auditory noise (low-noise or high-noise). This second factor allowed us to explore the neural mechanisms supporting reliability-weighted integration of auditory and visual signals (Magnotti, Ma, & Beauchamp, 2013). As shown in **Figure 1**, subjects were presented with words (either auditory-alone or AV) one at a time and were required to identify, via a button press, the initial consonant heard from four options. For example, on a trial with the auditory word ‘buy’, the options presented to the participant are the letters ‘b’, ‘g’, ‘d’, and ‘th’, which correspond to the words ‘buy’, ‘guy’, ‘die’, and ‘thigh’. When the incongruent visual stimulus ‘guy’ is paired with the auditory stimulus ‘buy’, participants report fusion percepts (‘d’ or ‘th’ consonants) at higher frequency due to the McGurk effect. To balance the presentation of AV pairings and to investigate incongruities that typically do not generate fusion percepts, we included matched “non-McGurk” trials (e.g., auditory ‘guy’ and ‘visual ‘buy’). This trial type does not elicit fusion responses but still reduces accuracy and slows reaction time.

**Figure 1.**
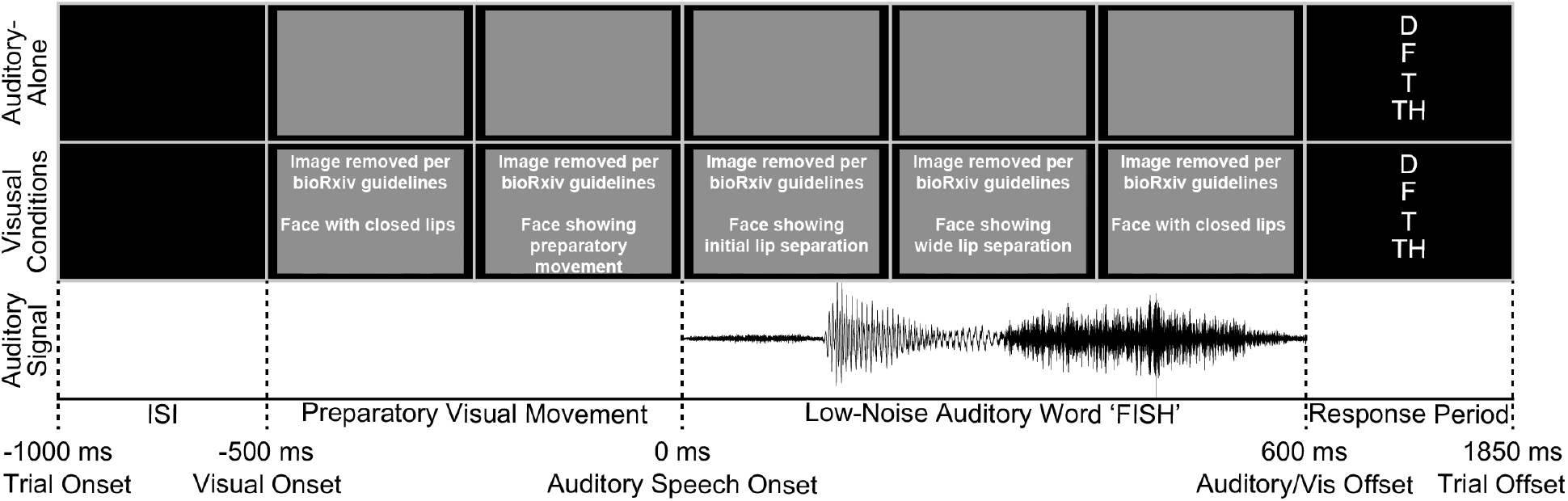
Trial schematic showing the auditory and visual stimuli for the word ‘fish’. All trials began with a blank screen during the ISI (random duration of 375 - 625 ms). Audio signals were muted until the first consonantal burst related to the word. 500 ms prior to sound onset, either a grey screen (auditory-alone condition) or a video (visual conditions) appeared. Auditory words were presented as recorded (low-noise conditions; SNR = 32.2 dB) or with pink-noise mixed into the signal (high-noise conditions; SNR = 6.4 dB). Following auditory/visual offset, participants identified which initial speech sound they heard from four options.

The paradigm included 40 trials per main condition (auditory-alone, congruent, incongruent) with half of the 40 words used in each condition presented in high-noise or low-noise contexts (counter-balanced across participants and testing sessions). In the low-noise conditions, the auditory stimuli were presented as they were recorded. In the high-noise condition, pink noise was added to reduce the SNR of the signal to −6 dB SPL. In the original paper (Ross et al., 2007), this level of noise was associated with approximately 55% accuracy in identifying the correct word in the auditory-alone condition (whereas accuracy approached 85% in the low-noise condition).

The order of trials was randomized for each participant. In the auditory-alone condition, a grey rectangle (same size and mean luminance of the videos) was presented 500 ms before sound onset to provide the same temporal cues to trial onset as in the visual conditions. Stimuli offset occurred 600 ms following sound onset. Non-clinical control participants’ task included a visual-alone condition that was not included in patients’ task, so was excluded from analyses.

Differing from the original study’s usage of free-response, we presented participants with four options (reflecting the initial consonant of the spoken word) to reduce difficulty for a clinical population and to allow for more reliable measures of reaction time. Participants were provided 1.25 seconds to respond, at which time the trial would auto-advance (missed trials were counted as incorrect). After each auto-advanced trial, the maximum wait time would increase by .25 seconds to a maximum of 2 seconds. Conversely, once participants responded before the maximum wait time for 10 trials in a row, this wait time decreased in .25 second intervals to a minimum value of 1.25 seconds.

#### Longitudinal Testing

To measure whether AV mechanisms can recover after damage, patients completed the AV speech paradigm at multiple time-points at clinically scheduled follow-up appointments. The median number of sessions completed was 5 (range = 2 - 9 sessions) over a median of 396 days (range = 95 to 792 days). To examine test-retest reliability in a non-patient population, 43 of the original 50 undergraduate control participants completed a second session, 3 - 13 days after the initial session (mean = 6.2 SD = 2.1).

### Quantification and Statistical Analyses

Accuracy and RT data were examined across patients with lesions extending into the left pSTS relative to patients with tumors in the frontal lobe as a control, as this region is sufficiently far from the pSTS to limit distal effects of a tumor and as a previous patient with Broca’s Aphasia demonstrated a preserved McGurk effect (Andersen & Starrfelt, 2015). Data were analyzed using a 3 x 2 repeated measures ANOVA with factors of visual-type (no visual [auditory-alone], incongruent, congruent) and auditory noise-level (low-noise, high-noise). Although the original degrees of freedom are reported here for clarity, p values were subjected to Greenhouse–Geisser correction where appropriate (Greenhouse & Geisser, 1959).

To examine functional recovery over time, we compared behavioral data as a function of condition and session across the groups. AV usage was calculated as the difference in accuracy between the congruent and incongruent conditions at each session. Then, for each participant, a linear regression was fit to these data across all testing sessions (reflecting the change in AV usage over time). Regression slopes were compared between patients with lesions in the pSTS and those with lesions in the frontal lobe. To examine whether patients with pSTS damage showed altered performance, we computed z-scores relative to frontal tumor patients on measures of overall accuracy for the proportion of fusion responses made during the AV Incongruent McGurk conditions. Z-scores were transformed into one-tailed p-values based on the expectation of weaker AV integration.

## RESULTS

### Non-Clinical Control Group

Session 1 results from *n* = 50 undergraduate participants are shown in Figure 2a-b. Accuracy data (Figure 2a) revealed main effects of visual-type [*F*(2,98) = 307.1, *p* = 1.77E-30, *η_p_^2^* = .862], noise-level [*F*(1,49) = 392.1, *p* = 4.98E-25, *η_p_^2^* = .889], and a significant interaction between the two [*F*(2,98) = 158.5, *p* = 4.35E-27, *η_p_^2^* = .764]. Reaction time data (Figure 2b) mirror those of the accuracy data, with main effects of visual-type [*F*(2,98) = 100.7, *p* = 3.03E-22, *η_p_^2^* = 0.673], noise-level [*F*(1,49) = 134.8, *p* = 1.12E-15, *η_p_^2^* = 0.733], and a significant interaction between the two [*F*(2,98) = 4.59, *p* = 0.015, *η_p_^2^* = .086]. Results were inconsistent with a speed-accuracy tradeoff. For example, under high noise, congruent visual information both improved accuracy [*t*(49) = 7.67, *p* = 6.17E-10, *d* = 1.08] and sped responses [*t*(49) = 6.2418, *p* = 9.91E-08, *d* = 0.883] relative to the auditory-alone condition. The test-retest reliability for accuracy on AV Incongruent trials was highly stable between session 1 and 2 (Figure 2C) [*r*(41) = .544, *p* = 1.63E-04] (*n*=43 completed the retest session), justifying this metric for an individual differences approach and for longitudinal testing in patients.

**Figure 2.**
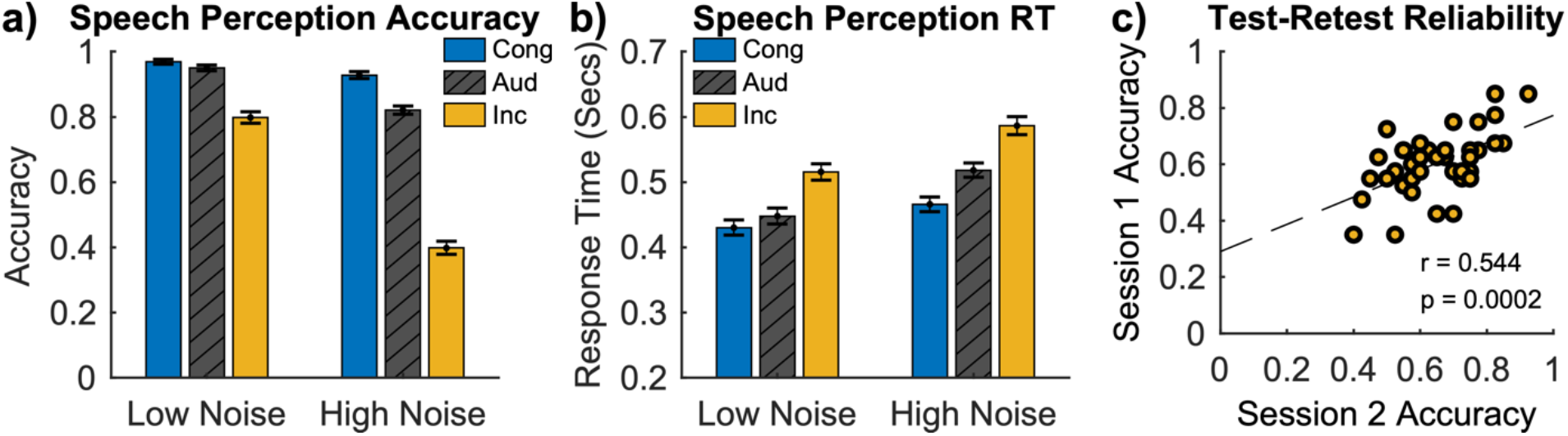
Normative data from control participants’ first testing session. (a) Accuracy in the main experimental conditions from the AV speech paradigm (Congruent AV, Auditory-Alone, Incongruent AV in low- and high-levels of auditory noise). Data show that high noise reduced auditory accuracy but also resulted in greater usage of visual information (leading to improved performance when the visual signal was congruent and reduced performance when incongruent). (b) Response time data for the same conditions presented in panel a. Error-bars reflect SEM. (c) Test-retest reliability for accuracy in the Incongruent AV condition at two time-points. Lower accuracy indicates greater illusion during McGurk trials.

Syllables that result in an experience of the McGurk effect can have an asymmetrical relationship. For example, an auditory /ba/ paired with a visual /ga/ typically leads to a McGurk fusion percept such as /da/, whereas an auditory /ga/ paired with a visual /ba/ typically will not lead to a fusion percept but, rather, a perceived /ba/ (auditory), /ga/ (visual), or /bga/ (combination) (MacDonald & McGurk, 1978; McGurk & MacDonald, 1976). In order to balance all stimuli across conditions, half of the Incongruent AV trials used combinations that should fail to produce the McGurk effect, but should still yield multisensory conflict (and possibly increased rates that the visual stimulus is ‘heard’). To examine these effects, we separated trials into those with stimulus combinations that lead to the McGurk effect versus those that do not (Figure 3). Consistent with prior research, we confirmed that participants reported a significantly greater number of fusion percepts (purple bars in Figure 3, reflecting response options 3 and 4) for McGurk stimuli compared to Non-McGurk stimuli [*t*(49) = 12.0, *p* = 3.50E-16, *d* = 1.70], and a significantly greater number of visual stimuli being reported as being heard (green bars in Figure 3, reflecting response option 2) for Non-McGurk stimuli compared to McGurk stimuli [*t*(49) = 12.5, *p* = 7.08E-17, *d* = 1.77].

**Figure 3.**
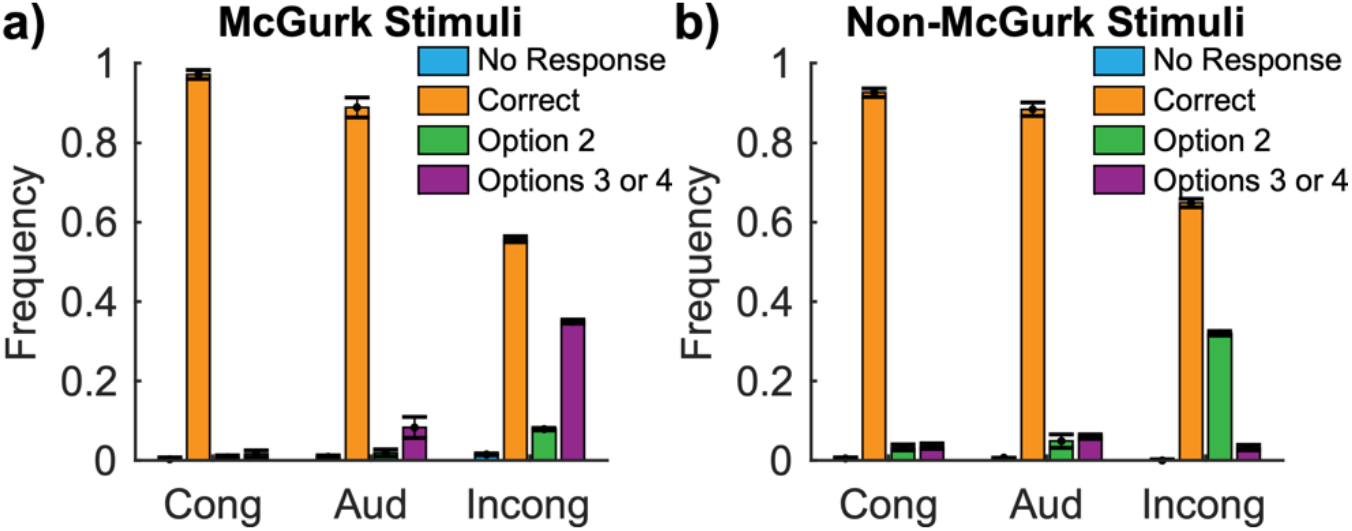
Normative data from control participants’ first testing session. Data averaged across low and high noise conditions, showing the frequency of response-types heard in the auditory-alone condition, or when paired with congruent or incongruent visual signals during trials either utilizing the (a) ‘McGurk’, or (b) ‘Non-McGurk’ stimuli. Previous research has demonstrated that the stimuli used in (a) are likely to create McGurk fusion percepts (e.g., auditory ‘Bill’ paired with visual ‘Gill’ will often lead to the experience of ‘Til’ or ‘Dill’), whereas the stimuli in (b) fail to evoke fusion percepts (e.g., auditory ‘Gill’ paired with visual ‘Bill’). The 4 response options were selected for each trial based on the incongruent condition so that they included the auditory stimulus (Correct), the visual stimulus (Option 2), or fusion percepts that indicate an experience of the McGurk effect (Options 3 and 4). The same response options were presented in each condition to enable relevant comparisons. Error-bars reflect SEM.

### Frontal Tumor Patients

Patients with an intrinsic brain tumor located in the frontal lobe (*n* = 14) showed a similar pattern of results to those of healthy controls during their initial testing session (prior to surgical removal of their tumor) (Figure 4). Accuracy data revealed main effects of visual-type [*F*(2,26) = 107.7, *p* = 5.06E-9, *η_p_^2^* = 0.892] and noise-level [*F*(1,13) = 105.4, *p* = 1.32E-7, *η_p_^2^* = 0.890], as well as a significant interaction between the two [*F*(2,26) = 44.8, *p* = 0.000001, *η_p_^2^* = 0.775]. Reaction time data mirror those of the accuracy data, with main effects of visual-type [*F*(2,26) = 32.9, *p* = 7.94E-7, *η_p_^2^* = 0.717], noise-level [*F*(1,13) = 23.4, *p* = 0.0003, *η_p_^2^* = 0.643], and a significant interaction between the two [*F*(2,26) = 5.68, *p* = 0.015, *η_p_^2^* = 0.304]. Results were inconsistent with a speed-accuracy tradeoff. For example, under high noise, congruent visual information both improved accuracy [*t*(13) = 9.99, *p* = 1.81E-07, *d* = 2.67] and sped responses [*t*(13) = 5.87, *p* = 5.50E-05, *d* = 1.61] relative to the auditory-alone condition. As in the healthy controls, the test-retest reliability for accuracy on AV Incongruent trials was high between the pre-Op and the initial post-Op session (Figure 4C) [*r*(8) = .731, *p* = .016].

**Figure 4.**
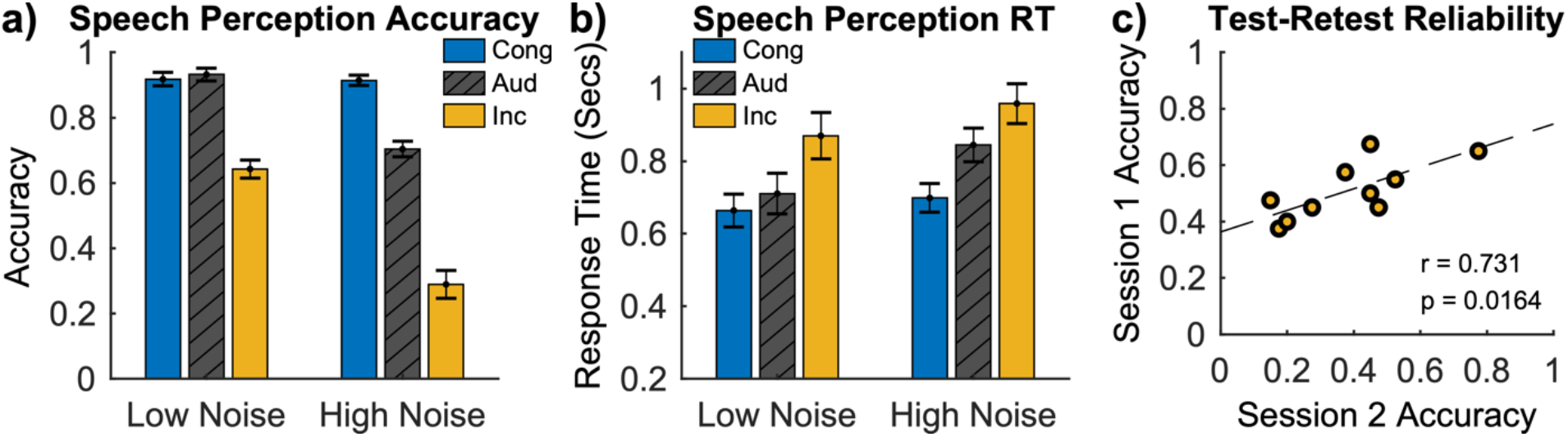
Patients with an intrinsic brain tumor in the frontal lobe showed a similar pattern of results compared to healthy controls (shown in Figure 2). Session 1 accuracy (a) and response time (b) data are shown from the main experimental conditions. Error-bars reflect SEM. (d) Test-retest reliability for accuracy in the Incongruent AV condition at the pre-Op and post-Op sessions.

Comparing accuracy between frontal tumor patients and healthy controls across all conditions showed a large main effect of group [*F*(1,62) = 29.8, *p* = 8.85E-7, *η_p_^2^* = 0.325], with healthy controls demonstrating greater accuracy in general. To more appropriately examine the impact of visual information on speech perception accuracy between these groups, we matched an equal number of controls to frontal patients based on their performance on the high-noise auditory-alone condition (as opposed to the low-noise condition to avoid ceiling effects) and conducted paired t-tests. Across these matched groups, no significant differences in accuracy were observed for either high-noise congruent AV trials [*t*(13) = 0.787, *p* = 0.4455, *d* = 0.210] or high-noise incongruent AV trials [*t*(13) = 1.459, *p* = 0.1683, *d* = 0.390]. Indeed, frontal tumor patients showed numerically *greater benefits* in the congruent condition and *worse performance* in the incongruent condition, consistent with prior research demonstrating that the presence of a frontal lesion does not reduce the integration of AV speech information (Andersen & Starrfelt, 2015).

As in control participants, to examine whether the pattern of response options chosen differed across conditions, we next separated trials into those with stimuli combinations that lead to the McGurk effect versus those that do not (Figure 5). Consistent with results from control participants, frontal tumor patients reported a significantly greater number of fusion experiences (purple bars in Figure 5, reflecting response options 3 and 4) for McGurk stimuli compared to Non-McGurk stimuli [*t*(13) = 9.13, *p* = 5.13E-07, *d* = 2.44], and a significantly greater number of visual stimuli being reported as being heard (green bars in Figure 5, reflecting response option 2) for Non-McGurk stimuli compared to McGurk stimuli [*t*(13) = 8.28, *p* = 1.53E-06, *d* = 2.21].

**Figure 5.**
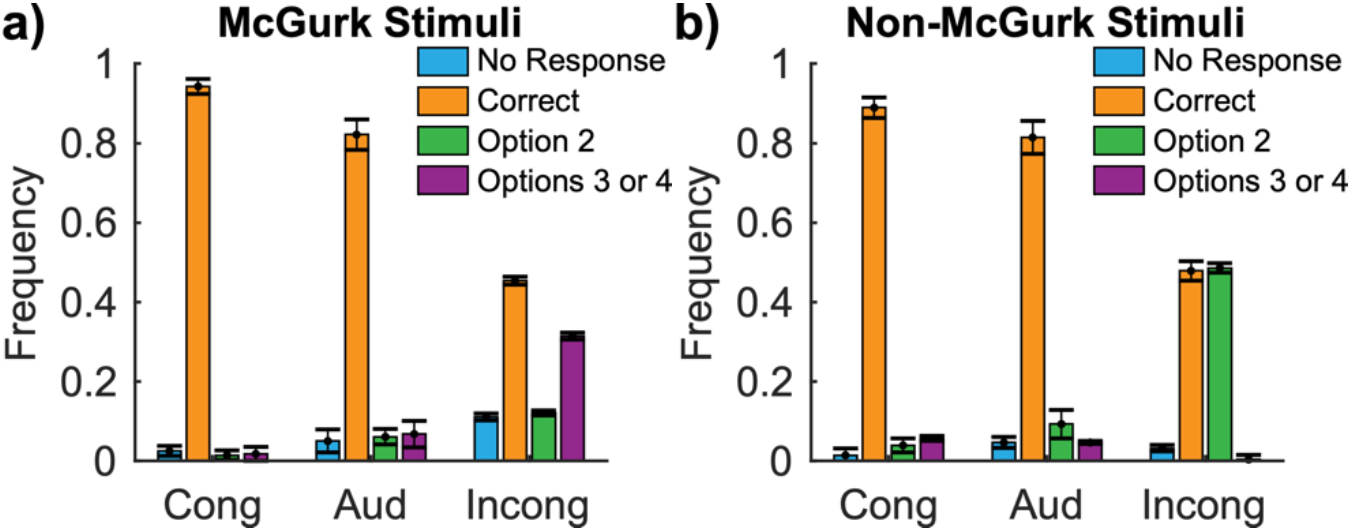
Data from patients with an intrinsic brain tumor in the frontal lobe averaged across low and high noise conditions, showing the frequency of response-types heard in the auditory-alone condition, or when paired with congruent or incongruent visual signals during trials either utilizing the (a) ‘McGurk’, or (b) ‘Non-McGurk’ stimuli. Error-bars reflect SEM.

### Patient 0036SF

Patient 0036SF had a WHO Grade 2 intrinsic brain tumor located at the boundary of the occipital, temporal, and parietal lobes, adjacent to the left pSTS (Figure 6). During pre-operative testing they showed no deficits on standard clinical measures of language performance. To examine their overall performance on the AV speech paradigm, we conducted a repeated measures ANOVA (random effect of session), examining their performance across their first three testing sessions (7 days prior to surgery, and then 2 and 84 days after surgery). Across these three sessions, accuracy data (Figure 6a) showed a very different pattern of results relative to healthy controls and frontal tumor patients, with no significant main effects of visual-type [*F*(2,4) = 1.000, *p* = 0.424, *η_p_^2^* = 0.333] or noise-level [*F*(1,2) = 0.0, *p* = 1.0, *η_p_^2^* = 0.0], nor did they show a significant interaction between the two [*F*(2,4) = 0.731, *p* = 0.536, *η_p_^2^* = 0.268]. Reaction time data showed a similar pattern, with no main effects of visual-type [*F*(2,4) = 0.724, *p* = 0.486, *η_p_^2^* = 0.266] or of noise-level [*F*(1,2) = 0.014, *p* = 0.917, *η_p_^2^* = 0.007], and no interaction between the two [*F*(2,4) = 2.26, *p* = 0.265, *η_p_^2^* = 0.530]. Indeed, examining the simplest comparison as in the other groups, under high noise, congruent visual information neither improved accuracy [*t*(2) = 0.654, *p* = 0.580, *d* = 0.378] nor sped responses [*t*(2) = 0.428, *p* = 0.710, *d* = 0.247] relative to the auditory-alone condition.

**Figure 6.**
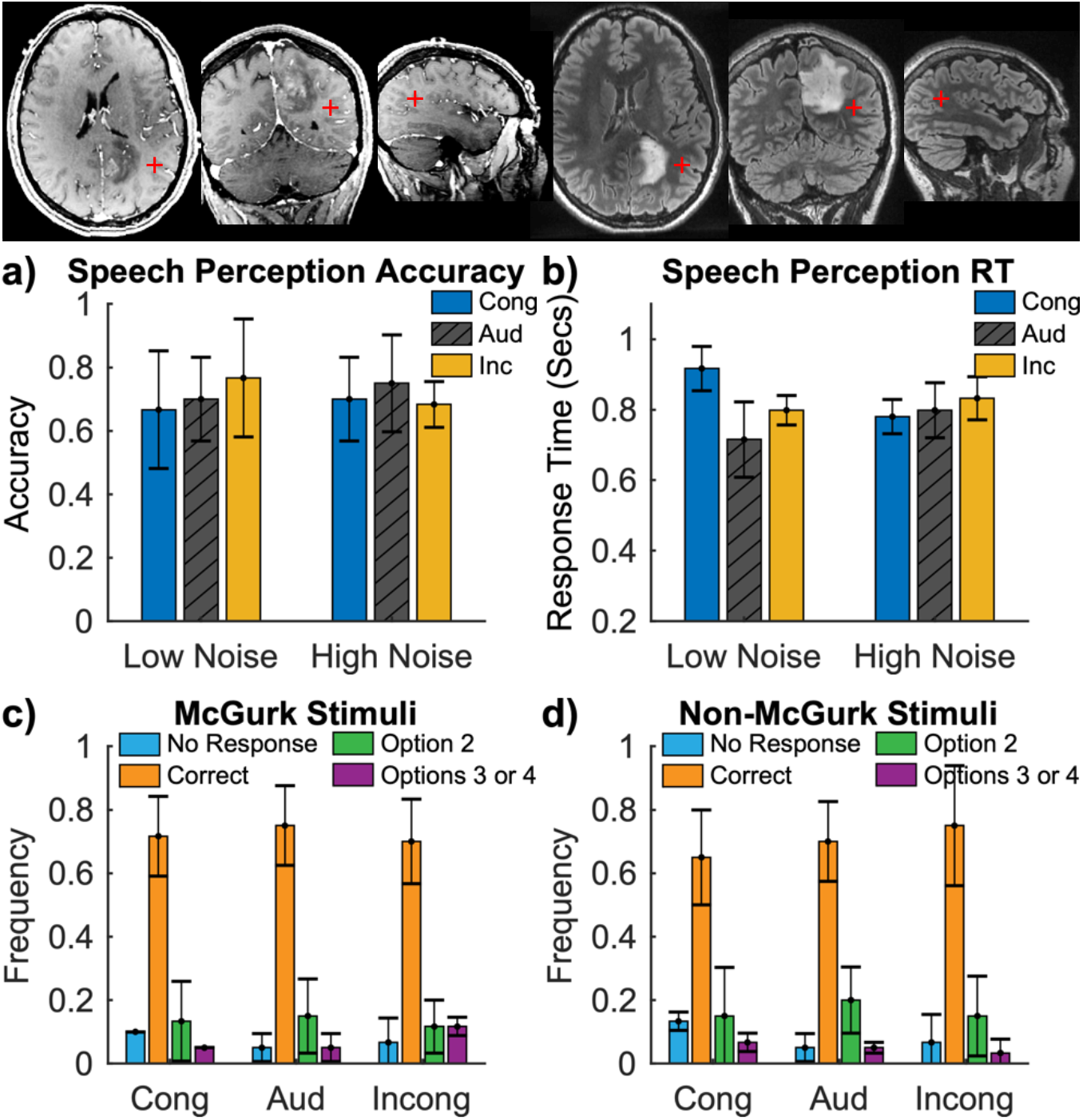
Data from a patient with an intrinsic brain tumor neighboring the left pSTS (shown by red plus sign), averaged across their initial 3 sessions. This individual showed reduced AV integration behaviors relative to frontal tumor patients, including no benefit of congruent visual information, either in terms of accuracy (panel a) or RT (panel b), and reduced influences of incongruent visual information. Similarly, the patient showed reduced ‘fusion’ responses during McGurk trials and fewer visual responses during Non-McGurk trials relative to frontal tumor patients. Top row of images shows the tumor on a post-gadolinium T1 (left) and Flair (right).

To examine whether the pattern of responses differed across conditions, we separated trials into those with stimuli combinations that lead to the McGurk effect versus those that do not (Figure 6c-d). Differing from control participants and frontal tumor patients, 0036SF demonstrated no difference in the number of fusion experiences (purple bars in Figure 6c, reflecting response options 3 and 4) for McGurk stimuli compared to Non-McGurk stimuli [*t*(2) = 2.50, *p* = .1296, *d* = 1.44], nor any difference in the number of visual stimuli reported as heard (green bars in Figure 6d, reflecting response option 2) for Non-McGurk stimuli compared to McGurk stimuli [*t*(2) = 1.00, *p* = .4226, *d* = .5774].

The between-session data for 0036SF indicated no usage of visual information; however, analyses based on this patient’s data alone may be weakly powered. To test whether their AV integration behaviors were indeed reduced relative to patients with a tumor in frontal regions, we next compared accuracy, responses, and response times across the groups. Compared to the frontal tumor group, 0036SF showed less benefit of congruent visual information relative to auditory-alone trials (*Z* = 1.738, *p* = .041) and less of a negative impact of incongruent visual information relative to auditory-alone trials (*Z* = 2.648, *p* = .004) (i.e., this patient was more accurate on incongruent AV trials than the frontal tumor group). Examining the impact of visual information on response times, 0036SF demonstrated significantly less benefit on their response times for congruent versus auditory-alone trials than frontal tumor patients (*Z* = 2.182, *p* = .015) and no difference relative to frontal tumor patients between incongruent and auditory-alone trials (*Z* = .557, *p* = .289). Finally, consistent with the view that 0036SF demonstrated reduced AV integration behaviors, they also showed a trend towards fewer ‘fusion’ responses on AV Incongruent McGurk trials compared to frontal tumor patients in terms of both the simple number of fusion responses made (*Z* = 1.467, *p* = .071) and in comparison to the number of ‘fusion’ responses made in AV Incongruent non-McGurk trials vs. AV Incongruent McGurk trials (*Z* = 1.538, *p* = .062).

### Patient 0012SF

Patient 0012SF had a WHO Grade 4 intrinsic brain tumor with a necrotic core located in the left pSTS and edema in neighboring areas (Figure 7). During pre-operative testing they showed no deficits on standard clinical measures of language performance. To examine their overall performance on the AV speech paradigm, we conducted a repeated measures ANOVA (random effect of session), examining their performance across their first three testing sessions (1 day prior to surgery, and then 7 and 89 days after surgery). In contrast to patient 0036SF, across these three sessions, accuracy data (Figure 7a) showed a similar pattern of results to those of healthy controls and frontal tumor patients, with significant main effects of visual-type [*F*(2,4) = 34.156, *p* = 0.0273, *η_p_^2^* = 0.945] and noise-level [*F*(1,2) = 41.3, *p* = 0.0234, *η_p_^2^* = 0.954], and a significant interaction between the two [*F*(2,4) = 43.0, *p* = 0.0225, *η_p_^2^* = 0.956]. Reaction time data mirrored those of the accuracy data, with a significant main effect of visual-type [*F*(2,4) = 34.292, *p* = 0.006, *η_p_^2^* = 0.945], a marginal main effect of noise-level [*F*(1,2) 6.860, *p* = 0.120, *η_p_^2^* = 0.774], and a marginal interaction between the two [*F*(2,4) = 12.186, *p* = 0.072, *η_p_^2^* = 0.859]. Results were inconsistent with a speed-accuracy tradeoff. For example, under high noise, congruent visual information both improved accuracy [*t*(2) = 7.56, *p* = 0.0171, *d* = 4.36] and sped responses [*t*(2) = 6.515, *p* = 0.0228, *d* = 3.762] relative to the auditory-alone condition.

**Figure 7.**
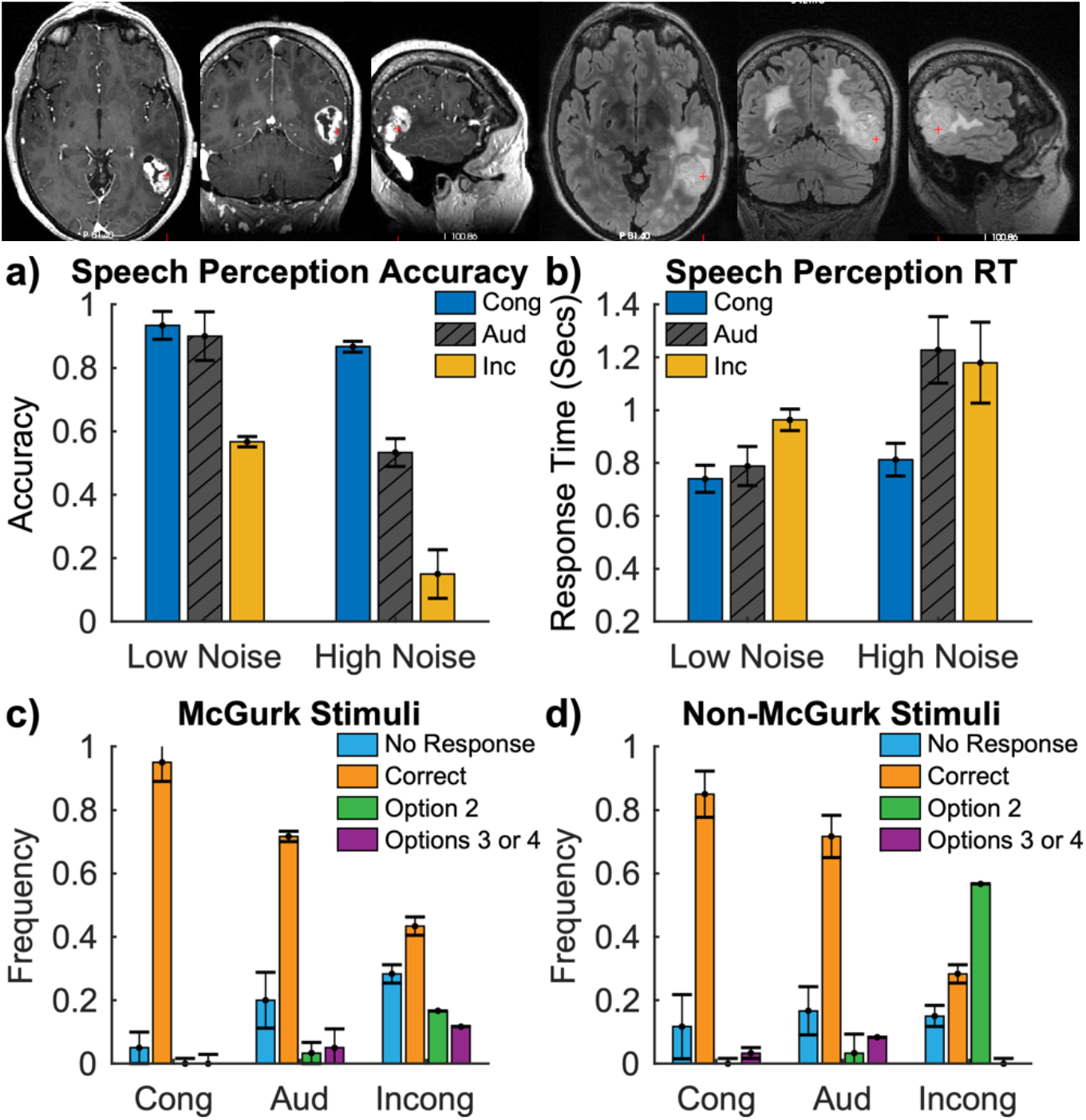
Data from a patient with an intrinsic brain tumor in the left pSTS, averaged across their initial 3 session, and who demonstrated a similar pattern of behaviors to both healthy controls and frontal tumor patients. Critically, in spite of left pSTS damage, the patient showed intact benefits of congruent visual information, both in terms of accuracy (panel a) and RT (panel c), as well as some impairments due to incongruent information. However, during McGurk trials (panel b) but not non-McGurk trials (panel d) the patient showed significantly fewer fusion responses that normally indicate an experience of the illusion. Top row of images shows the tumor on a post-gadolinium T1 (left) and Flair (right).

To examine whether the pattern of responses differed across conditions, we separated trials into those with stimuli combinations that lead to the McGurk effect versus those that do not (Figure 7c-d). Differing from control participants and frontal tumor patients, but consistent with the results of the other left pSTS patient, 0012SF demonstrated no difference in the number of fusion experiences (purple bars in Figure 6c, reflecting response options 3 and 4) for McGurk stimuli compared to Non-McGurk stimuli [*t*(2) = 1.94, *p* = .1917, *d* = 1.121], but did show a significant difference in the number of visual stimuli reported as heard (green bars in Figure 7d, reflecting response option 2) for Non-McGurk stimuli compared to McGurk stimuli [*t*(2) = 13.9, *p* = .0052, *d* = 8.00].

Patient 0012SF’s between-session data indicated strong benefits of congruent visual information on speech perception, but critically, no McGurk effect. We next sought to examine whether this usage was reduced relative to patients with a tumor in frontal regions as was observed in 0036SF through the comparison of accuracy, response choice, and response times across the groups. Compared to the frontal tumor group, 0012SF showed no difference in their usage of congruent visual information relative to auditory-alone trials (*Z* = 1.20, *p* = .116) and no difference in the negative impact of incongruent visual information relative to auditory-alone trials (Z = 0.020, p = .492). Indeed, this patient showed a numerically larger benefit of congruent visual information on speech perception compared to the frontal tumor group, in contrast to the results of 0036SF and predictions that left pSTS damage should impair all AV speech integration behaviors. Examining the impact of visual information on response times, 0012SF demonstrated a trend towards *greater* benefit for congruent versus auditory-alone trials compared to frontal tumor patients (*Z* = 1.62, *p* = .052) (i.e., congruent visual information led to greater response time speeding than observed in frontal tumor patients) and no difference relative to frontal tumor patients between incongruent and auditory-alone trials (*Z* = 0.491, *p* = .312). Finally, in contrast to the ‘typical’ pattern of results shown for congruent AV stimuli, patient 0012SF showed trends towards fewer ‘fusion’ responses on AV Incongruent McGurk trials compared to frontal tumor patients in terms of the simple number of fusion responses made (*Z* = 1.47, *p* = .071), and a trend in the same direction for the comparison to the number of ‘fusion’ responses made in AV Incongruent non-McGurk trials vs. AV Incongruent McGurk trials (*Z* = 1.33, *p* = .092).

### Longitudinally Tracking of AV Speech Perception Performance

Both the frontal tumor patients and the two patients with left pSTS tumors were tested on this paradigm at multiple time-points over a two-year period to examine whether impaired AV speech integration behaviors would recover over time. Figure 8 shows longitudinal accuracy data for all tumor patients in each of the main three experimental conditions. Frontal tumor patients (Figure 8a) and 0012SF (Figure 8b) each showed large influences of visual information on speech perception accuracy (both improvements from congruent visual signals and costs from incongruent visual signals) that were highly stable over time. In contrast, patient 0036SF (Figure 8c) initially showed no influence of visual information, but at 270 days following tumor resection, began to demonstrate benefits both from congruent visual signals and costs from incongruent visual signals. Note that 0036SF showed a general drop at their initial post-operative testing session, which reflects general cognitive impairment following surgery that is present in some patients.

**Figure 8.**
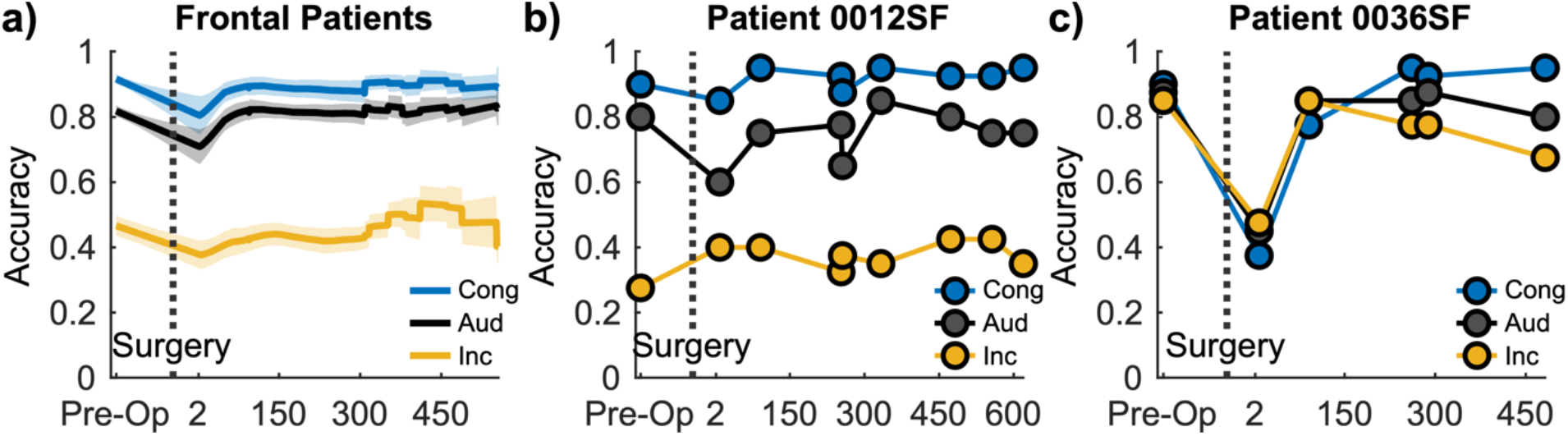
Longitudinal performance in frontal tumor patients (left), patient 0012SF (middle), and patient 0036SF (right). Data show accuracy in each of the three main experimental conditions at different testing sessions; y-axis reflects the date of testing relative to the pre-op session. Tumor removal occurred 1-2 days following the initial pre-op session. (a) Mean performance of frontal tumor patients over time (shading reflects SE) showing stable benefits from congruent visual information and costs from incongruent visual information. (b) Patient 0012SF showed a similarly stable and robust pattern of results relative to frontal tumor patients, despite having 3 resections in the left pSTS over this two-year period. (c) Patient 0036 initially showed no influence of visual information as they were equally accurate in each of the three conditions. However, beginning ~270 days following their resection they showed increasingly large multisensory effects (such that congruent visual information improved performance while incongruent visual information impaired it), consistent with functional recovery through neural plasticity.

To quantify their usage of visual information, we conducted a repeated measures ANOVA (random effect of session), examining 0036SF’s performance across their final three testing sessions (Figure 9). While their initial testing sessions showed no evidence of visual usage, their final sessions better matched the results of the other patients, with significant main effects of visual-type [*F*(2,4) = 24.0, *p* = 0.030, *η_p_^2^* = 0.923] and noise-level [*F*(1,2) = 306.3, *p* = 0.003, *η_p_^2^* = 0.994], and a marginal interaction between the two [*F*(2,4) = 7.14, *p* = 0.102, *η_p_^2^* = 0.781]. Reaction time data mirrored this pattern, with a marginal main effect of visual-type [*F*(2,4) = 12.9, *p* = 0.065, *η_p_^2^* = 0.866], a significant main effect of noise-level [*F*(1,2) 28.4, *p* = 0.033, *η_p_^2^* = 0.934], and no interaction between the two [*F*(2,4) = 1.73, *p* = 0.318, *η_p_^2^* = 0.464]. Results were inconsistent with a speed-accuracy tradeoff. For example, under high noise, congruent visual information both improved accuracy [*t*(2) = 4.91, *p* = 0.039, *d* = 2.84] and non-significantly sped responses [*t*(2) = 2.27, *p* = 0.151, *d* = 1.312] relative to the auditory-alone condition. Indeed, compared to the frontal tumor group, 0036SF’s final testing session showed no significant difference in their usage of congruent visual information relative to auditory-alone trials (*Z* = 1.17, *p* = .120) (numerically a larger difference than frontal patients) and a smaller, but still significantly smaller difference in the negative impact of incongruent visual information relative to auditory-alone trials (Z = 1.82, p = .034). Particularly noteworthy, 0036SF’s recovery was most pronounced for AV stimuli in high noise auditory contexts, potentially indicating a strategically learned behavior utilized only when necessary (e.g., when auditory signals are significantly degraded).

**Figure 9:**
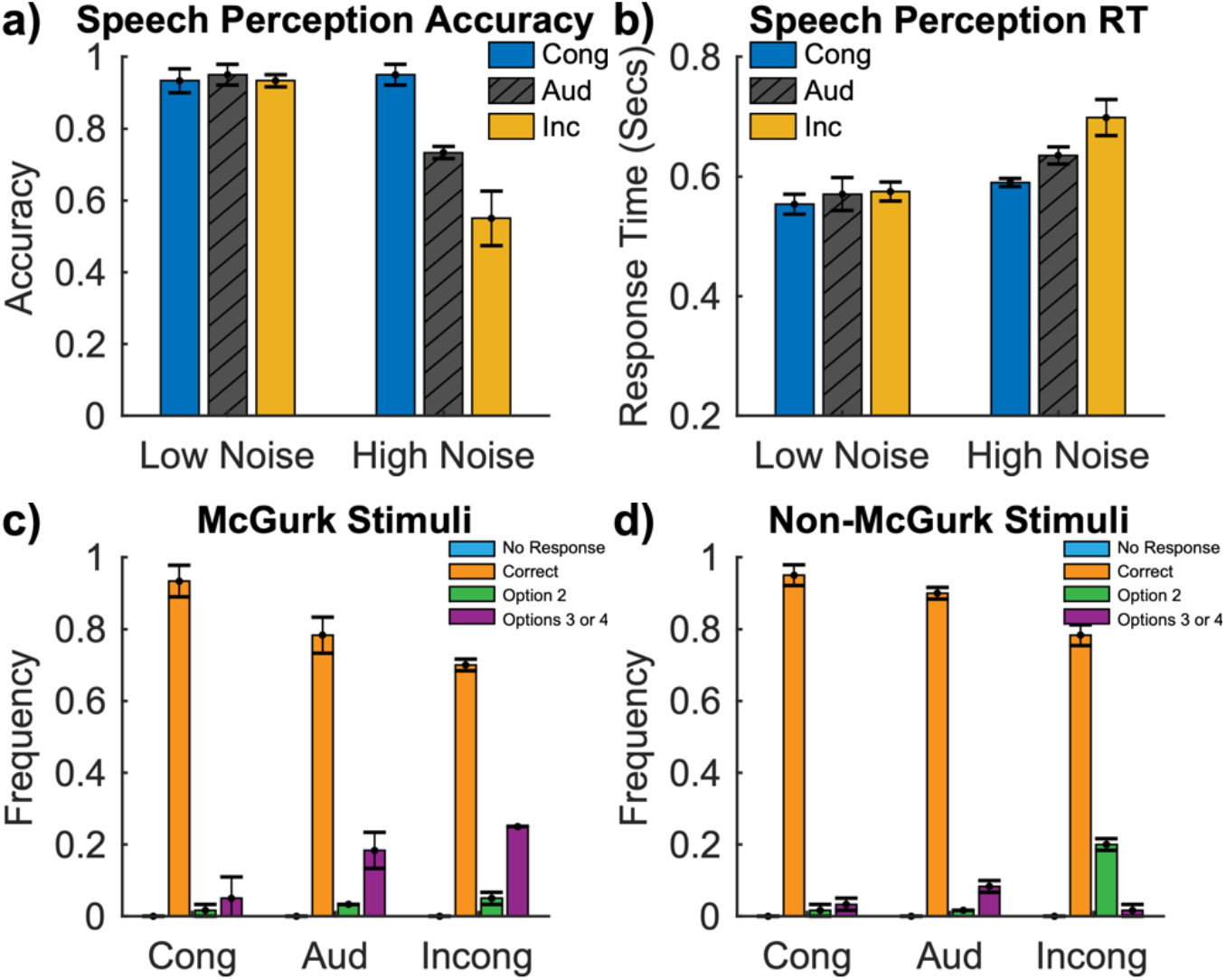
Data from patient 0036SF, averaged across their final 3 sessions. After 9 months of demonstrated AV impairments, this individual showed improved AV integration behaviors, improved accuracy and faster RTs to congruent AV stimuli, and lower accuracy and slower RTs to incongruent AV stimuli. Additionally, this patient showed a stronger McGurk effect as denoted by the number of fusion percepts reported. Effects of visual information were maximal in high noise conditions. Errorbars reflect SEM.

Finally, to more directly examine whether the strength of the McGurk effect varied over time in these individuals, we calculated the proportion of fusion responses made for Incongruent AV McGurk trials. Frontal tumor patients showed a stable proportion of McGurk fusion responses over time (mean = 32.5%). Comparing the final session across patients, 0036SF no longer showed a difference relative to frontal tumor patients (*Z* = .020, *p* = .492), while 0012SF showed a trend towards a persistent reduction in the number of fusion responses (*Z* = 1.41, *p* = .080).

## Discussion

Our novel paradigm uses ecologically valid stimuli (whole words as opposed to single phonemes) to quantify the influence of visual information on multiple aspects of speech perception, eliciting robust and reliable effects in both typically developing individuals and patients with intrinsic brain tumors. Broadly, these data demonstrate that congruent visual signals facilitate speech perception, whereas incongruent visual signals impair or modulate it. Additionally, these data confirm that noise significantly affects the strength of visual influences on speech perception. When auditory noise is elevated (rendering the auditory signal less reliable), participants use visual information more, leading to improved performance when the visual signals are congruent and reduced performance when the signals are incongruent.

The present work centers on two patients with damage to the left pSTS who showed different patterns of behavior and divergent patterns of recovery following surgery. Little research to date has sought to address the causal role of the left pSTS in AV speech perception. Using non-invasive transcranial magnetic stimulation (TMS), Beauchamp and colleagues (Beauchamp et al., 2010) demonstrated that inhibitory TMS applied over the left pSTS, but not a control site, reduced the strength of the McGurk effect. However, as TMS was applied at only one experimental site, it is unclear if other regions exist that contribute to AV speech integration processes. Indeed, prior research has demonstrated that preparatory lip movements can modulate auditory processes through direct projections from visual motion area MT/V5 to auditory cortex, effectively bypassing the pSTS (Besle et al., 2008). Similar reports of a weakened McGurk effect have been reported in patients following strokes near the left pSTS (Hickok et al., 2018). However, the McGurk effect may reflect only a very specific form of AV speech integration (conscious extraction of viseme information) and may be of limited relevance to other forms of AV integration including visual timing information (Besle et al., 2008; Chandrasekaran, Trubanova, Stillittano, Caplier, & Ghazanfar, 2009), spectral recovery through lip shape (John Plass, Brang, Suzuki, & Grabowecky, 2020), and speaker identity information (Brang, 2019; Vatakis & Spence, 2007).

Consistent with these findings, we found that a tumor in the left pSTS resulted in a weaker McGurk effect (in terms of the number of fusions experienced) compared to patients with a tumor in the frontal lobe. However, one of the individuals (0012SF) persisted in showing preserved signs of robust auditory-visual speech facilitation (with accuracy and response time benefits to congruent visual signals) in spite of a high-grade glioma centered in the left pSTS. This demonstration provides a critical dissociation between facilitative and modulatory AV speech functions and is consistent with models suggesting that the McGurk effect and facilitative AV speech benefits likely rely on partly different neural mechanisms (Arnal et al., 2009; Arnal et al., 2011).

While damage to the left pSTS can lead to a reduction in both the benefits of visual signals on speech perception and the McGurk effect, critically, our longitudinal data suggest that these behaviors can recover over time in some cases. This is consistent with a prior case report showing a McGurk effect within the normal range of the typically developing population in a patient tested 5 years after a stroke in the left pSTS (Baum, Martin, Hamilton, & Beauchamp, 2012). Using fMRI, the authors demonstrated a larger recruitment of neural activity in the right pSTS, consistent with a view that functional plasticity allowed a persistent experience of the McGurk effect. However, as this patient was not tested longitudinally, it is possible that this patient never showed any deficit due to the stroke. Moreover, this study did not examine more ecologically valid benefits of congruent visual signals on speech perception as demonstrated here.

In light of this evidence, we predict that the right pSTS will show structural changes as behavior recovers, consistent with research demonstrating that homologous structures in the non-lesioned hemisphere compensate for neural damage during language processing (Dronkers, Wilkins, Van Valin Jr, Redfern, & Jaeger, 2004) and demonstrate increased contralesional gray matter volume during aphasia recovery (Xing et al., 2016). The observation that AV speech benefits can recover after their loss is important because multisensory behaviors are critical to helping impaired sensory processing. Patients who show auditory deficits along with AV speech deficits are likely to have a worse long-term outcome compared to those with auditory deficits alone, especially if remnant multisensory interactions can serve as a scaffolding to retrain auditory speech perception abilities.

Notably, however, while 0012SF showed robust AV facilitations across nine testing sessions (over ~2 years), they failed to show any recovery of the McGurk effect, in contrast with patient 0036SF. One possible explanation is that 0012SF had a total of three resections in this region during this time-period, which may have limited the extent to which functional plasticity engaged or its resistance to repeated damage. Alternatively, as 0012SF showed robust benefits of AV congruity effects throughout the entire time-course, there may have been little reason for the brain to recover the processes that enable the McGurk effect (i.e., the McGurk effect as a spandrel).

One potential concern in the current study is that neural plasticity may occur in patients with an intrinsic brain tumor before surgical resection. Specifically, we cannot exclude the possibility that the preserved AV speech integration behaviors demonstrated in 0012SF were not due to already recovered behaviors. Three pieces of evidence lead us to disfavor this view. First, high-grade gliomas typically produce little neural plasticity prior to resection due to their fast-growing nature, which leads to greater deficits upon their resection (Desmurget, Bonnetblanc, & Duffau, 2007; Duffau, 2005). Second, the recovery of language functions takes a significant amount of time, with clinical recovery of language functions typically plateauing within 6-9 months following damage (Pedersen, Stig Jørgensen, Nakayama, Raaschou, & Olsen, 1995). Third, this patient showed preserved AV speech integration behaviors in spite of an impaired McGurk effect.

These data emphasize that AV speech integration is not a unitary phenomenon dependent on a single multisensory hub in the left pSTS and that the McGurk effect does not capture all AV speech behaviors. Indeed, it is possible that prior research leading to an overemphasis on the importance of the pSTS was due to broad usage of the McGurk effect as a representative manipulation. In contrast, our data are more consistent with converging behavioral and neurophysiological evidence suggesting that auditory enhancements from congruent visual speech (e.g., better detection and faster reaction times) and visual modulations of what is heard (i.e., the McGurk effect) are subserved by two distinct mechanisms (Arnal et al., 2009; Arnal et al., 2011; Baart et al., 2014; K. Eskelund, J. Tuomainen, & T. S. Andersen, 2011).

This distinction may reflect a neural dissociation between predictive multisensory interactions that optimize feedforward encoding of auditory information and later feedback processes that alter auditory representations via the pSTS (Arnal et al., 2009; Arnal et al., 2011). In support of this view, both visual speech (Arnal et al., 2009; Arnal et al., 2011; Besle, Fort, Delpuech, & Giard, 2004; van Wassenhove, Grant, & Poeppel, 2005) and other anticipatory visual cues (Vroomen & Stekelenburg, 2009) can speed and reduce the magnitude of early physiological responses associated with auditory feedforward processing, potentially reflecting optimization of auditory encoding in accordance with temporal or acoustic constraints imposed by visual information. These early feedforward effects, which are insensitive to AV congruity in speech, are temporally, topologically, and spectrally distinct from later (>300 ms) responses that are specific to crossmodally incongruent speech (Arnal et al., 2009; Arnal et al., 2011; van Wassenhove et al., 2005). These later incongruity-specific interactions point to a hierarchical feedback regime in which unisensory speech processing is altered in accordance with integrated AV information from the pSTS (Arnal et al., 2011; Kayser & Logothetis, 2009; Olasagasti, Bouton, & Giraud, 2015).

In summary, these data demonstrate that while damage to the left pSTS is associated with a weaker McGurk effect, it does not universally reduce the ecologically important benefits of congruent visual information on speech perception. This suggests a dissociation in neural mechanisms such that the pSTS reflects only one of multiple critical areas necessary for AV speech interactions. Moreover, when neural damage does affect AV speech integration, patients can recover these behaviors over time, which will in turn enable better long-term resistance to speech perception deficits.

## Acknowledgements

This study was supported by NIH Grants R00DC013828, F32DC018199, and K08NS110919.

